# Genome sequencing, *de novo* assembly and annotation of the commercially important bamboo, *Bambusa tulda* Roxb

**DOI:** 10.1101/2025.07.19.665512

**Authors:** Sutrisha Kundu, Oliver Rupp, Sonali Dey, Mridushree Basak, Sudeshna Bera, Annette Becker, Malay Das

## Abstract

*Bambusa tulda* Roxb., a member of the Bambusoideae subfamily, is an ecologically and commercially important plant resource widely distributed on the Indian subcontinent. Our study reports long-read PacBio HiFi sequencing and genome assembly of *B. tulda*. Flow cytometry analysis revealed its estimated genome size ∼3 Gb. The *de novo* genome assembly of *B. tulda* predicted 43 contigs, distributed across three subgenomes, with a total size of 1.37 Gb, contig N50 of 35.69 Mb, and BUSCO score 99%. Repetitive elements constitute 63.31% of the genome. Functional annotation predicted 56,890 protein-coding genes, constituting 19.44% of the genome. This high-quality draft genome assembly will serve as an invaluable resource for future studies on the life history traits, phylogenomic analysis, comparative genomics, and targeted genome modification for important trait improvement of *B. tulda*.

## Background and summary

Bamboos are among the fastest growing, non-timber, renewable plant resources and are widely distributed throughout the tropics and subtropics [1,2]. They comprise the Bambusoideae subfamily of the monocotyledonous grass family Poaceae, and have significant ecological and economical value [3,4]. The Bambusoideae subfamily is classified into two clades: herbaceous bamboos and woody bamboos [5]. Herbaceous bamboos possess weakly developed rhizomes, lower lignocellulosic biomass, and exhibit annual or seasonal flowering cycles [6]. In contrast, the woody bamboos demonstrate rapid culm growth, high accumulation of lignocellulosic biomass, and delayed flowering subsequent upon lengthy vegetative phases upto 120 years [1,7–9]. Although, the first bamboo genome *Phyllostachys heterocycla* (=*P*. *edulis*) was sequenced more than a decade ago [10], major progress in bamboo genome sequencing was achieved more recently due to the introduction of long read sequencing technologies and advancement in algorithms for genome assembly and annotation [11]. Twelve species from the Bambusoideae subfamily, with diverse worldwide distribution, have been sequenced and their chromosome-level assemblies have been published (Fig. 1a) [2,5].

**Fig. 1.**
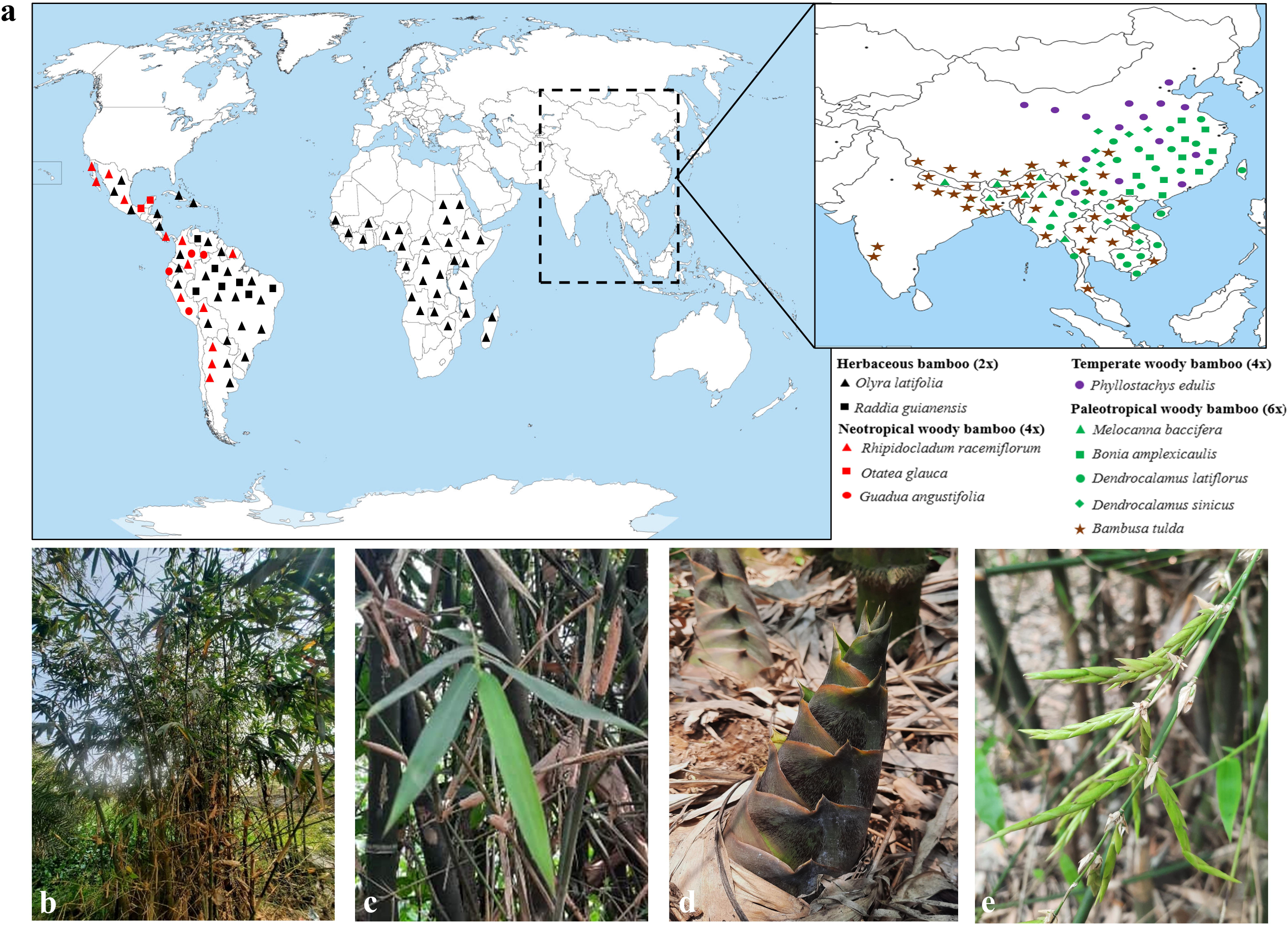
(a) Worldwide native distribution of sequenced bamboo genomes, alongwith *B. tulda*. The map was drawn based on information obtained from World Flora Online (WFO 2024), Kew (POWO, 2025), and Guadua Bamboo (https://www.guaduabamboo.com/). (b) *B. tulda* plant in native habitat, (c) leaf, (d) emerging culm, and (e) spikelet inflorescence.

*Bambusa tulda*, also known as the Bengal bamboo, is a paleotropical woody bamboo, native to the Indian subcontinent, and certain parts of southeast Asia (Fig. 1a, b, c, d, e). It is putatively aneuploid (70-72 chromosomes) [12] in nature, and possesses highly lignified culms. Its tentative flowering time is ∼50 years [13] and exhibits both sporadic and gregarious flowering [14]. In addition to their conventional use as construction wood in rural housing in south and southeast Asia, the plant draws renewed attention due to their potential use in the paper-pulp industry [15] and bio-energy sector [9]. *In vitro* micropropagation method have also been optimized for *B. tulda*, providing opportunity for genetic intervention [16]. Until now, some flowering-associated genes were identified for functional studies in *B. tulda* flowering [17–20]. However, in the absence of a good quality, whole genome sequence, comparative evolutionary studies and genetic manipulation for trait improvement remain challenging.

This study aimed to sequence, assemble, and annotate the *B. tulda* genome to provide a genomic resource for this valuable species. Long-read, PacBio high-fidelity (HiFi) sequencing was performed to obtain a draft genome assembly of *B. tulda*. The genome size estimated by flow-cytometry and *k*-mer analyses were ∼3 Gb and ∼1.17 Gb (-k 25, p: 2), respectively (Table 1, Fig. 2). Approximately 116.4 Gb raw PacBio HiFi data and 76.8 Gb transcriptome data was obtained (Tables 2, 3). The primary contig assembly, generated through HiFiasm, produced 752 contigs with the contig N50 value 31.02 Mb (Table 4). Two haplotypes were obtained from a haplotype-resolved assembly, with haplotype 1 containing 827 and haplotype 2 containing 326 contigs (Table 4). Further refinement and mapping of the primary assembly with other sequenced bamboo genomes produced a final draft genome assembly of 43 contigs with contig N50 value 35.69 Mb (Table 5). Genome assembly quality confirmed a BUSCO score of ∼99% (Fig. 3). Genome annotation identified 56,890 protein-coding genes, constituting 19.44% of the genome (Table 6). Repetitive sequences constituted major part (63.31%) of the *B. tulda* genome (Table 7). Additionally, 1,355 rRNA and 1019 tRNA genes were identified (Table 8). This draft genome assembly and annotation of *B. tulda* will serve as a valuable genetic resource for future studies (Box1).

**Fig. 2.**
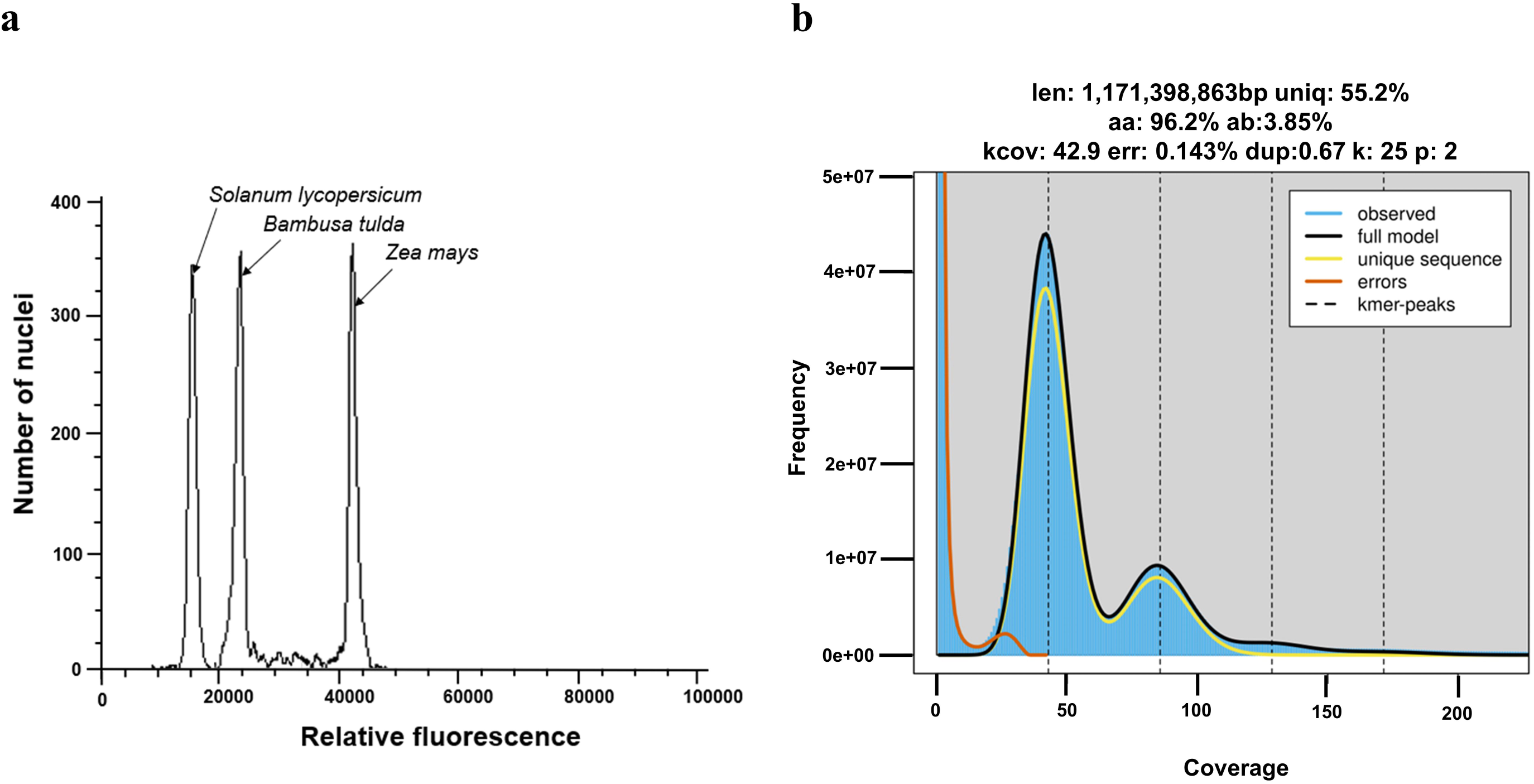
Genome size estimation of *B*. *tulda*. (a) Flow cytometry based estimation of genome size from fluorescence intensity histograms showing mean peak positions of *B. tulda*, *Solanum lycopersicum*, and *Zea mays*. (b) Genome size estimation based on *k*-mer distribution (“-k 25”) using Jellyfish-2 and Genomescope 2.0. The analysis also reveals unique sequences, with both homozygous and heterozygous distribution.

**Fig. 3.**
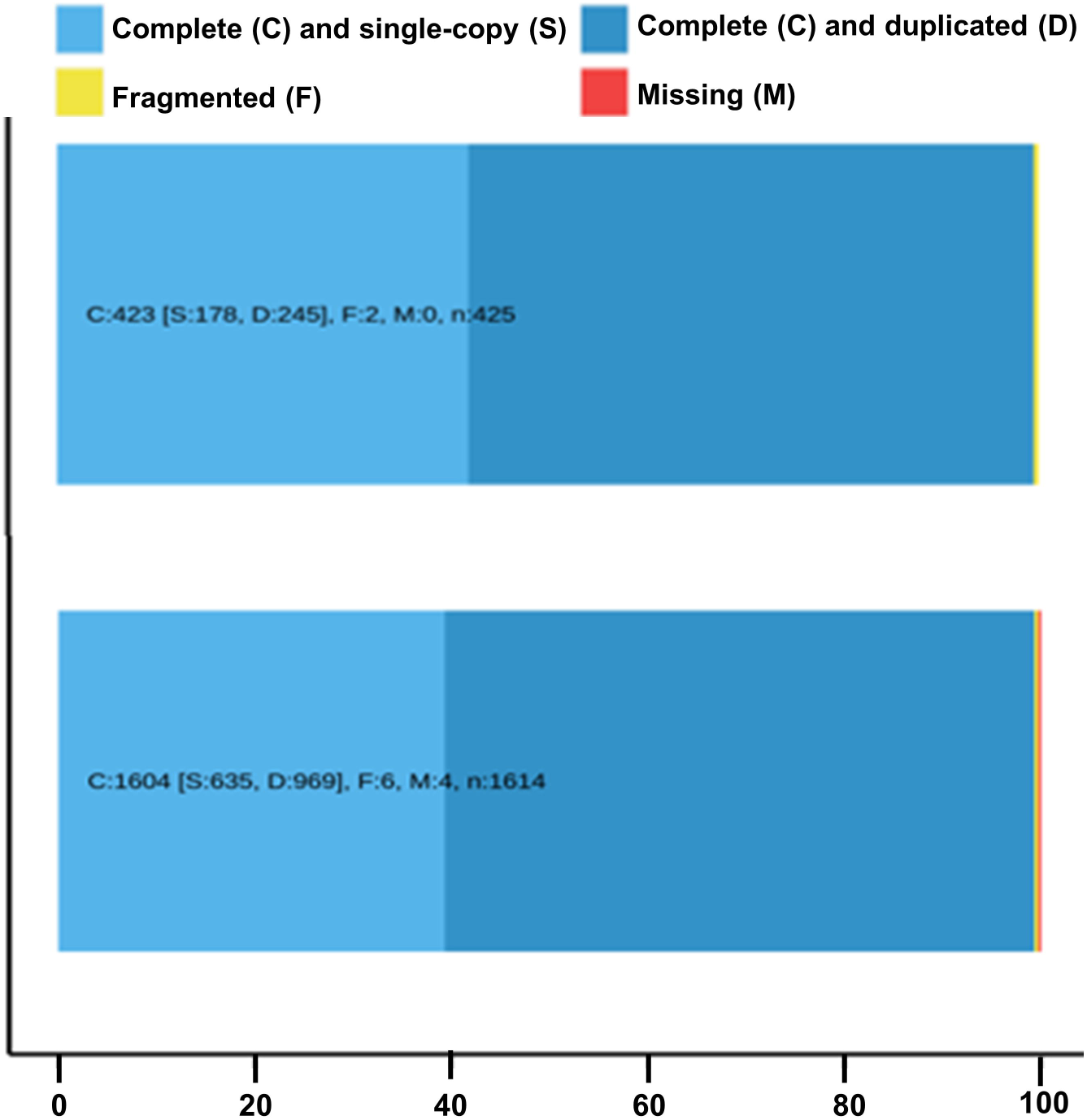
Evaluation of genome assembly completeness of *B. tulda* against the Embryophyte (n = 425) and Viridiplantae (n = 1614) lineage using BUSCO analysis, demonstrating the complete, fragmented, and missing BUSCO groups.

**Table 1.** Estimation of genome size using flow cytometry analyses.

| **Sample** | **Reference** | **G1 peak mean position of reference** | **G1 peak mean position of sample** | **Ratio** | **2C DNA content (pg)** | **Genome size (Gb)** |
| --- | --- | --- | --- | --- | --- | --- |
| *Bambusa tulda* | *Solanum lycopersicum* | 15557 | 24567 | 1.58 | 3.08 | 3.01 |
| *Bambusa tulda* | *Zea mays* | 42980 | 24328 | 0.57 | 3.09 | 3.02 |

**Table 2.** Sequencing statistics of *B. tulda* genome.

|  | **DNA1** | **DNA2** | **DNA1** | **DNA2** | **DNA3** | **Total** |
| --- | --- | --- | --- | --- | --- | --- |
| **Sequencing platform** | Pacio<br>Revio | PacBio<br>Revio | PacBio<br>Sequel IIe | PacBio<br>Sequel IIe | PacBio<br>Sequel IIe |  |
| **READS** | 5480695 | 1873037 | 1102778 | 734918 | 1072867 | 10264295 |
| **SIZE** | 54.99 gb | 24.64 gb | 13.75 gb | 9.20 gb | 13.80 gb | 116.38 gb |
| **MAX** | 40.10 kb | 43.91 kb | 35.37 kb | 38.96 kb | 38.55 kb | 43.91 kb |
| **MEAN** | 10.03 kb | 13.16 kb | 12.47 kb | 12.52 kb | 12.86 kb | 11.34 kb |
| **MEDIAN** | 10.21 kb | 12.72 kb | 12.10 kb | 12.46 kb | 12.37 kb | 11.38 kb |
| **MIN** | 111.00 bp | 89.00 bp | 102.00 bp | 97.00 bp | 138.00 bp | 89.00 bp |
| **N50** | 10.67 kb | 13.06 kb | 12.51 kb | 12.91 kb | 12.77 kb | 11.87 kb |

**Table 3.**
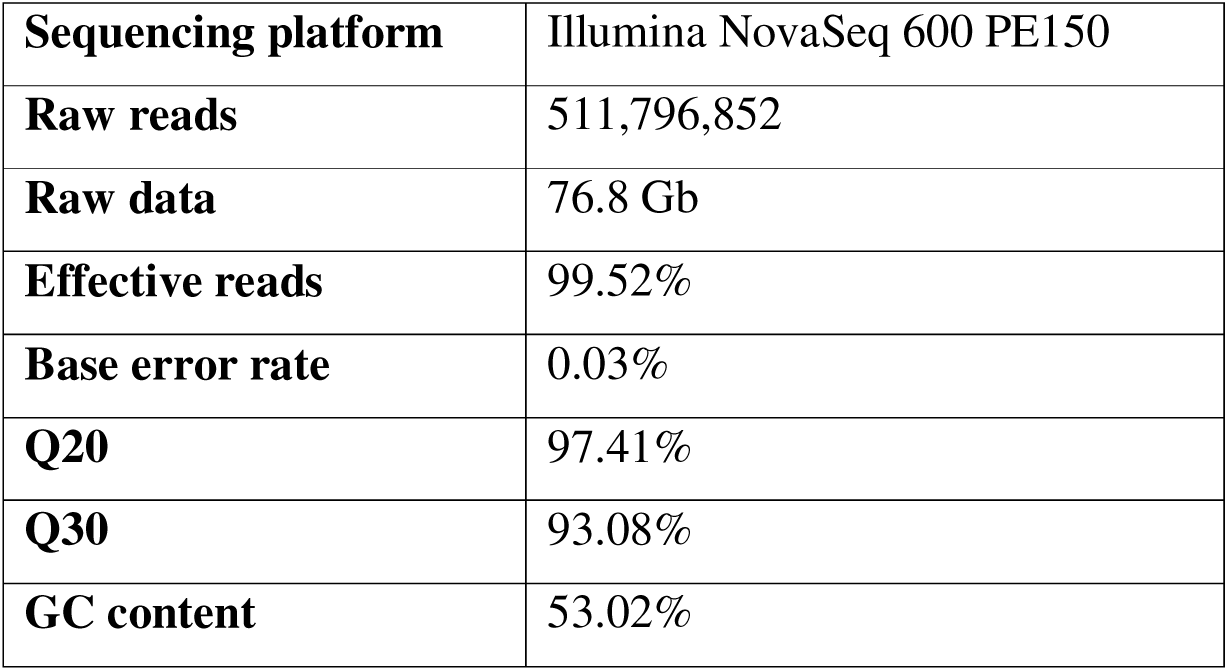
Sequencing statistics of *B. tulda* transcriptome.

|  |  |
| --- | --- |
| **Sequencing platform** | Illumina NovaSeq 600 PE150 |
| **Raw reads** | 511,796,852 |
| **Raw data** | 76.8 Gb |
| **Effective reads** | 99.52% |
| **Base error rate** | 0.03% |
| **Q20** | 97.41% |
| **Q30** | 93.08% |
| **GC content** | 53.02% |

**Table 4.** Assembly statistics of the primary and haplotype-resolved *B. tulda* genome.

|  | Primary assembly | Haplotype 1 | Haplotype 2 |
| --- | --- | --- | --- |
| **CONTIGS** | 752 | 827 | 326 |
| **SIZE** | 1.85 Mb | 1.38 Gb | 1.40 Gb |
| **MAX** | 63.43 Mb | 57.40 Mb | 63.43 Mb |
| **MEAN** | 2.46 Mb | 1.67 Mb | 4.30 Mb |
| **MEDIAN** | 30.58 kb | 30.15 kb | 49.41 kb |
| **MIN** | 4.40 kb | 4.40 kb | 15.70 kb |
| **N50** | 31.02 Mb | 26.77 Mb | 28.84 Mb |
| **L50** | 22 | 19 | 18 |
| **N90** | 15.11 Mb | 8.87 Mb | 10.98 Mb |
| **L90** | 52 | 50 | 47 |

**Table 5.** Assembly statistics of the final draft genome of *B. tulda*.

|  | Final draft assembly |
| --- | --- |
| **Contigs** | 43 |
| **Size** | 1.37 Gb |
| **Maximum** | 63.43 Mb |
| **Mean** | 31.77 Mb |
| **Median** | 30.48 Mb |
| **Minimum** | 10.51 Mb |
| **N50** | 35.69 Mb |
| **L50** | 15 |
| **GC content** | 44.61% |

**Table 6.** Statistics of gene prediction in *B. tulda* genome.

| Property | Value |
| --- | --- |
| Total gene length | 265,570,276 bp |
| Total gene number | 56890 |
| Genes percentage of Genome | 19.44% |
| Total Exon Number | 525887 |
| Average Exons per Gene | 5.88 |
| Total Exon Length | 177,027,591 bp |
| Exon Percentage of Genome | 12.96% |
| Total CDS Number | 485055 |
| Total CDS Length | 116,429,828 bp |
| CDS Percentage of Genome | 8.52% |
| Average Gene Length | 4668.14 bp |
| Average Exon Length | 336.63 bp |
| Average CDS Length | 240.03 bp |

**Table 7.**
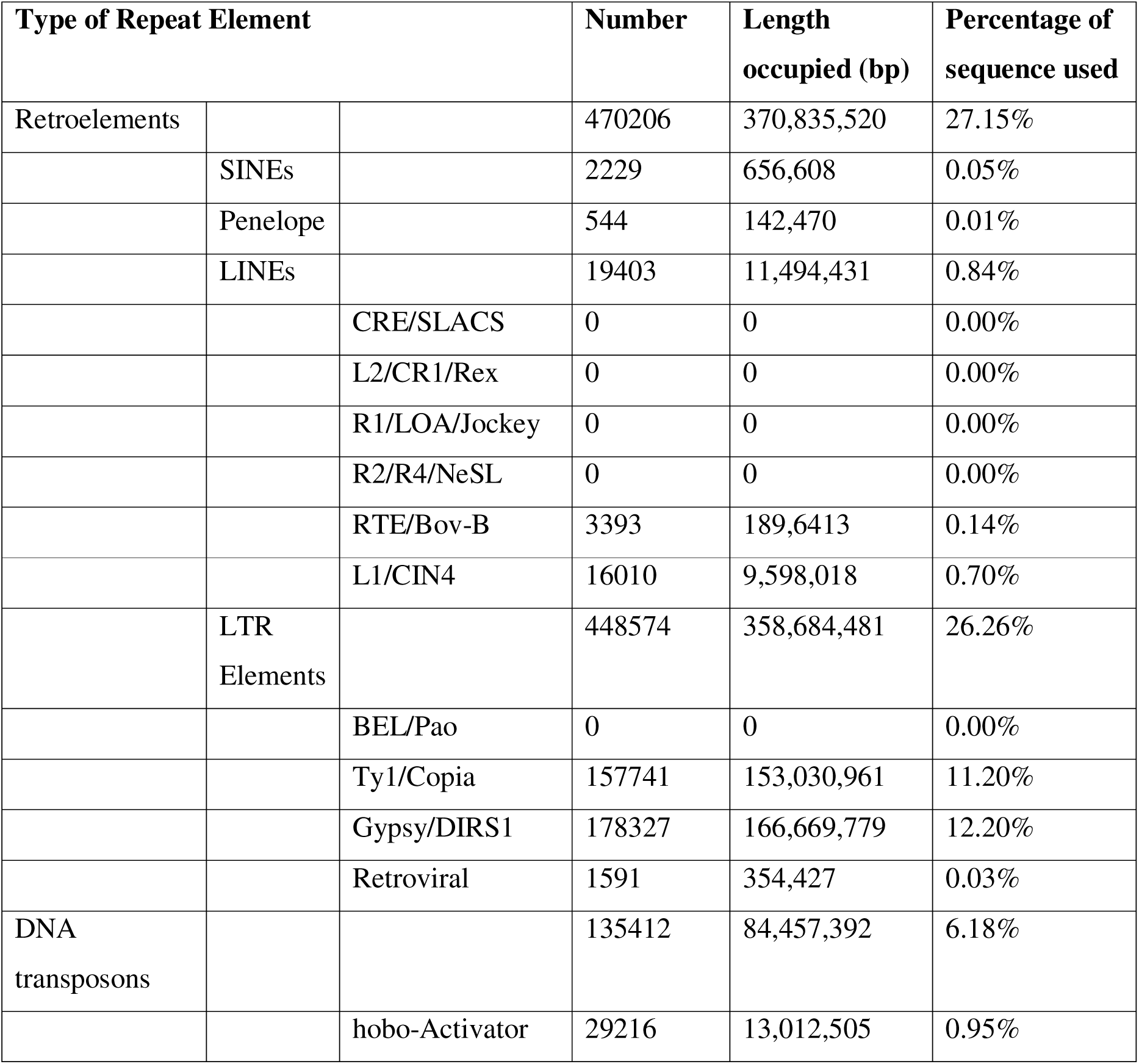
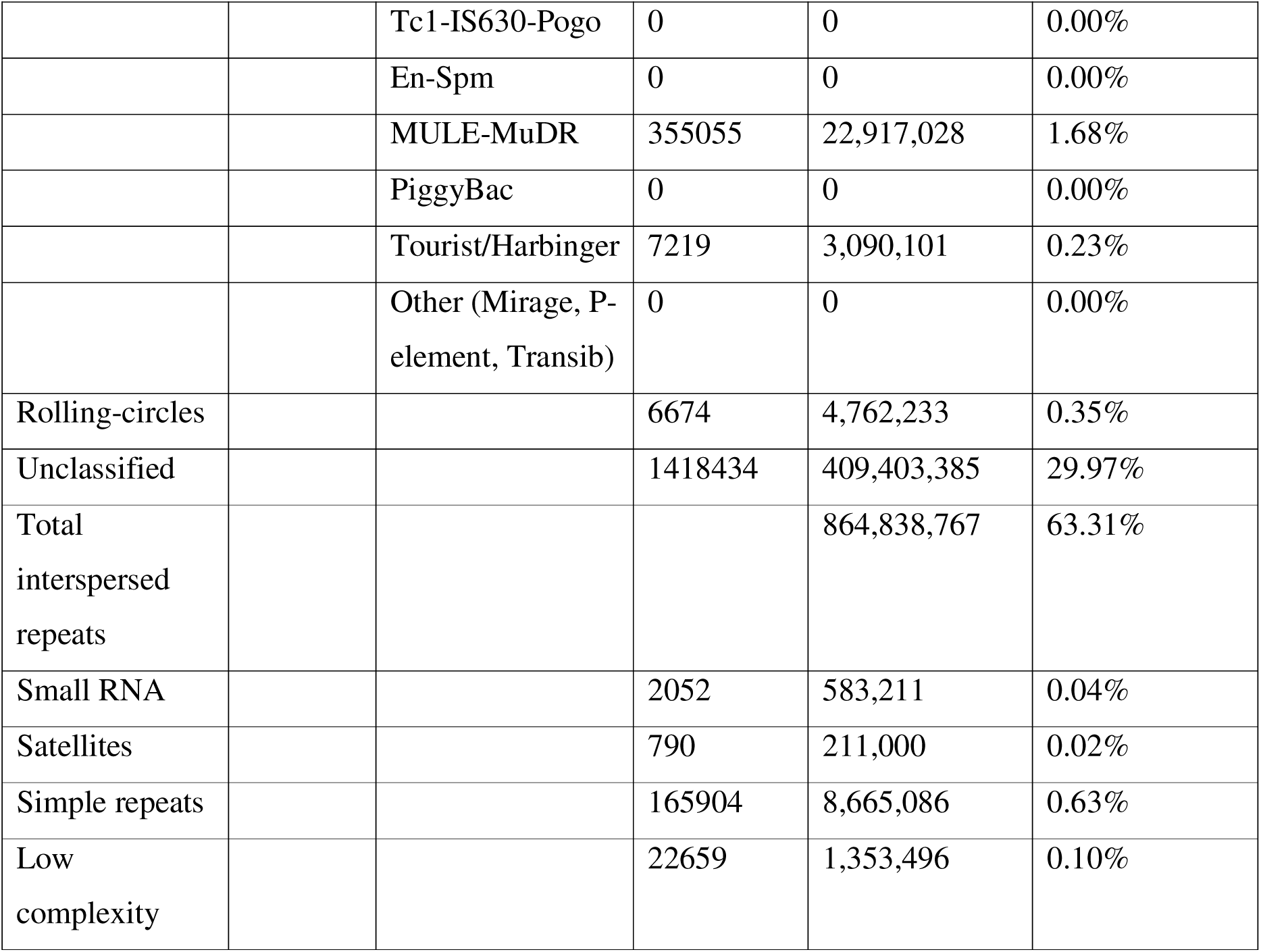
Statistics of repeat elements in the *B. tulda* genome.

**Table 8.** Statistics of non-coding rRNA and tRNA prediction in *B. tulda* genome.

| Type of ncRNA | Count | Total Length (bp) | Average Length (bp) | % of Genome |
| --- | --- | --- | --- | --- |
| 18S rRNA | 15 | 19703 | 1313.533333 | 0.001442 |
| 28S rRNA | 17 | 31624 | 1860.235294 | 0.002315 |
| 5.8S rRNA | 8 | 1223 | 152.875000 | 0.000090 |
| 5S rRNA | 1315 | 151008 | 114.834981 | 0.011054 |
| tRNA | 1019 | 77003 | 75.567223 | 0.005637 |

#### Box 1. Important research questions in *B. tulda* biology

- What is the polyploidization history of *B. tulda*?
- Why is flowering time delayed up to ∼50 years?
- What are the major regulators promoting rapid vegetative growth?
- What is the role of repeat elements in bamboo genome evolution?
- What are the major gene families that expanded/contracted in the bamboo lineages?

## Materials and methods

### Sample collection

Tissues were collected from a natural, flowering population of *B. tulda*, located at Rahuta, Shyamnagar (22.83°N, 88.40°E), West Bengal, India. Fresh young leaves collected were used for genome size estimation and whole genome sequencing, while for transcriptome sequencing, emerging culm was used (Figs. 1B, C, D). All the collected tissues were quickly flash-frozen in dry ice, and subsequently stored at -80°C.

### Genome size estimation by flow cytometry analysis

For genome size estimation, two plants with known genome sizes: *Solanum lycopersicum* L. Stupické polnı’ rané (1.96 pg) and *Zea mays* L. ‘CE-777’ (5.43 pg) were used as internal references. Isolation of nuclei was carried out by following the nuclei-isolation protocol of Dolezel [21]. In brief, 40 mg fresh young leaf tissue from both the reference plants and *B. tulda* were finely chopped in 1 ml ice-cold nuclei isolation Galbraith buffer. The homogenate was filtered through a 40 μm nylon mesh. The filtrate was treated simultaneously with 50 μg/ml RNase A and 50 μg/ml propidium iodide. After incubation in the dark for 30 mins, the suspension of stained nuclei was introduced into flow cytometer (S3e Cell Sorter, Bio-Rad), using argon-ion laser of 488 nm. Live gating was set around the fluorescence area signal (FL3-A) to obtain cell cycle histograms. Based on the fluorescence histograms, the nuclear DNA content was calculated. The estimated genome size of *B. tulda* was 3.01 Gb and 3.02 Gb on the basis of the two reference plants *S. lycopersicum* and *Z. mays*, respectively (Table 1).

### Genome and transcriptome sequencing

High molecular weight (HMW) genomic DNA was extracted from the young leaves using the NucleoBond® HMW DNA kit (Macherey-Nagel, Germany), following the manufacturer’s protocol. The extracted HMW genomic DNA was fragmented to obtain Single-Molecule Real-Time (SMRT) bell libraries. The SMRTbell libraries were then sequenced in the PacBio Sequel IIe (3 libraries) and PacBio Revio (2 libraries) platform (Novogene, Genomics Singapore Pte. Ltd) (Fig. S1, S2). The raw reads obtained by sequencing 5 libraries were pooled together and subjected to quality control. Genome sequencing in the PacBio Sequel IIe platform yielded 36.75 Gb data with N50 value of 12.73 kb and mean read length 12.62 kb (Table 2). Similarly, genome sequencing in the PacBio Revio platform yielded 79.63 Gb data with N50 value of 11.87 kb and mean read length 11.6 kb and (Table 2). From the two sequencing platforms, a total of 10264295 PacBio SMRT reads corresponding to 116.38 Gb data was obtained (Table 2).

For accurate annotation of the *B. tulda* genome, transcriptome sequencing was performed in the Illumina Novaseq 6000 platform. Total RNA was extracted from tissues using the RNeasy® Plant Mini Kit (Qiagen), following the manufacturer’s protocol. The integrity of isolated RNAs was examined in the Agilent 5400 Bioanalyzer, and sample with RNA integrity value >8 were used for library construction. The library was subjected to the Illumina NovaSeq 6000 PE150 platform for sequencing. Transcriptome sequencing yielded a total raw data output of 76.8 Gb with a Q30 value of 93.08% (Table 3).

### Estimation of genome characteristics

Following genome sequencing, *k*-mer analysis was performed to estimate the genome size, heterozygosity rate, and proportion of repetitive sequences of *B. tulda*. The distribution of *k*-mer frequency from the raw PacBio HiFi reads was performed in Jellyfish (v2.3.0) [22] software. The *k*-mer spectrum obtained for “-k 25”, ploidy = 2 was then analyzed using the GenomeScope 2.0 [23] web tool. The predicted genome size was ∼1.17 Gb, heterozygosity rate 3.85%, sequencing error rate 0.143% and repetitive sequences constituted 44.8% of the genome (Fig. 2). Smaller peaks at 3x and 4x coverage indicated possible genome duplication.

### *De novo* genome assembly

The clean PacBio HiFi reads were assembled *de novo* into a draft haplotype-resolved assembly and a primary contig assembly (a combination of both haplotypes) using Hifiasm v0.16.1-R341 [24] tool. The primary contig assembly produced a total of 752 contigs equivalent to 1.85 Gb of data with the N50 value 31.02 Mb (Table 4, Fig. S3a). Size of the two haplotype assemblies were 1.38 Gb for Haplotype 1 and 1.40 Gb for Haplotype 2 (Table 4, Fig. S3b). The assembled genome size was lower compared to that estimated by GenomeScope 2.0, possibly because of higher repetitive elements in the genome.

The 752 primary contigs obtained from the *B. tulda* HiFiasm assembly were further mapped and compared to other sequenced bamboo genomes. Finally, 43 individual contigs were obtained from the assembled draft genome (Fig. 4a). Among them, 39 contigs could be nomenclatured and divided into three distinct (A, B, and C) subgenomes based on their mapping and homology to chromosome-level assemblies of other bamboo genome. This indicated that *B. tulda* is possibly a hexaploid plant, where each chromosome is represented by three subgenomes. There are some duplicated contigs, which corresponded to a single chromosome in other bamboo genomes, but in this case, they could not be assembled together, and hence assigned as two different contigs. The remaining four contigs could not be assigned to regions of any specific chromosomes. The final draft genome assembly resulted in total genome size of 1.37 Gb, contig N50 value 35.69 Mb, mean contig length 31.77 Mb, and GC content was 44.61% (Table 5).

**Fig. 4.**
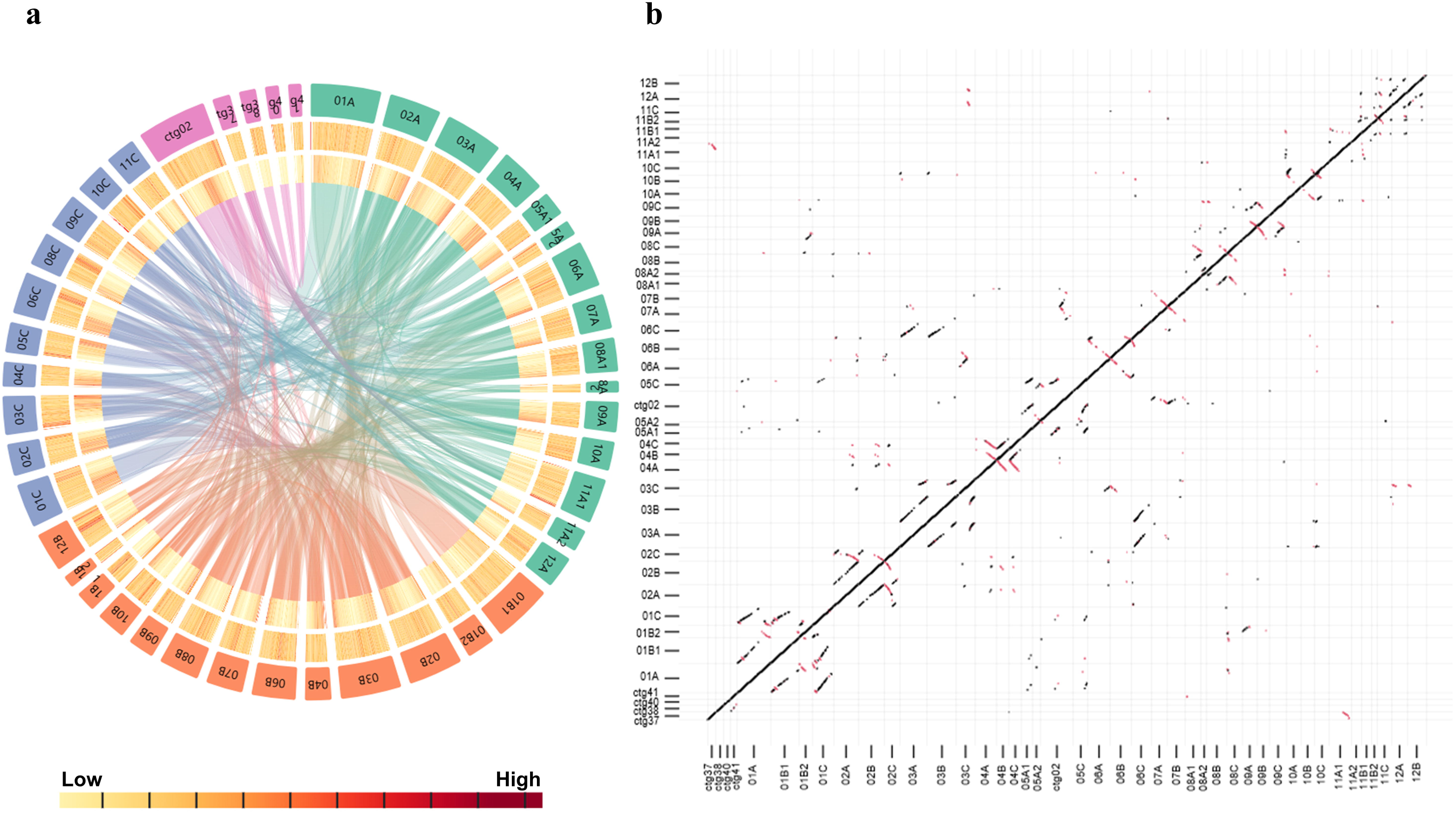
Genomic features for the 43 contigs of the *B. tulda* genome. (a) Circos plot representing the different genome features of the *B. tulda* genome assembly. The outer to inner tracks represent different contigs, GC content, transposable element density, and collinear blocks. Green, orange, and blue blocks represent contigs of subgenome A, subgenome B, and subgenome C, respectively. Purple blocks represent unidentified contigs. The plot was visualized using Accusyn software [50]. (b) Dot plot representing the subgenome syntenic relationship of *B. tulda* based on the annotated coding genes.

### Annotation of repetitive elements

To annotate repetitive elements in *B. tulda* genome, a combination of *de novo* and homology-based prediction was used. For *de novo* prediction, RepeatModeler [25,26] was used, which in turn uses two *ab initio* repeat sequence prediction softwares: RECON (v1.0.8) [27] and RepeatScout (v1.0.6)[28]. LTR_FINDER [29] was used specifically for *de novo* prediction of LTR elements. Homology-based method to identify the transposable elements involved using the RepeatMasker [30,31] software against Repbase database.

Approximately, 63.31% of the total *B. tulda* genome was annotated as repeat elements. The identified transposable elements were further classified into two broad categories: Class I retrotransposons and Class II DNA transposons. They represented 27.15% (370 MB) and 6.18% (84 MB) of repetitive sequences, respectively, while 29.97% repeat elements remained unclassified (Table 7, Fig. 4a).

### Gene prediction and functional annotation

For predicting protein-coding genes, a combination of three different methods were applied together: *de novo*, homology-based, and transcriptome-assisted. The *de novo* prediction was done using RNA-Seq and protein evidence data, allowing BRAKER3 [32] to perform automated genome annotation. GeMOMa v1.7 [33] was used for homology-based prediction, by mapping the previously sequenced bamboo gene models [5]. The transcriptome guided genome annotation was conducted using Trinity (v2.1.1) [34,35], HISAT2 (v2.2.1) [36], and StringTie (v1.3.3) [37,38]). The transcriptome data was mapped to the genome using HISAT2 to generate BAM files, which were then used as input for genome-guided transcriptome assembly in StringTie. Also, unigene assemblies were generated by *de novo* assembly of transcriptome data using Trinity and rnaSPADES [39]. The genes predicted from all three above methods were consolidated together using Evidence Modeler (EVM) v2.1.0 [40] and PASA [41,42] to finally obtain non-redundant protein-coding gene structures. In total, 56,890 protein-coding genes constituting 19.44% of the *B. tulda* genome were identified (Table 6). The predicted genes of *B. tulda* were mapped back to the *B. tulda* genome using GeMoMa [33], with the “synteny checker” module enabled. The dot plot was created with the “synplot.r” script (k=3) included in GeMoMa on the reference gene table created by the GeMoMa run (Fig. 4b).

For functional annotation, homology-based annotation was done using SwissProt and TrEMBL databases. The predicted protein-coding sequences were aligned against these two databases through BLASTP [43] analysis with e-value ≤1e^−5^. The functional domains of the proteins were identified using InterProScan [44].

### Annotation of rRNA and tRNA

Among the non-coding RNA genes, rRNAs and tRNAs were predicted using the *B. tulda* genome. The rRNA genes were predicted using Barrnap v0.9 [45], while the tRNA genes were predicted using tRNAscan-SE [46,47]. From this prediction, 1355 rRNA and 1019 tRNA genes were identified successfully (Table 8, Fig. 5).

**Fig. 5.**
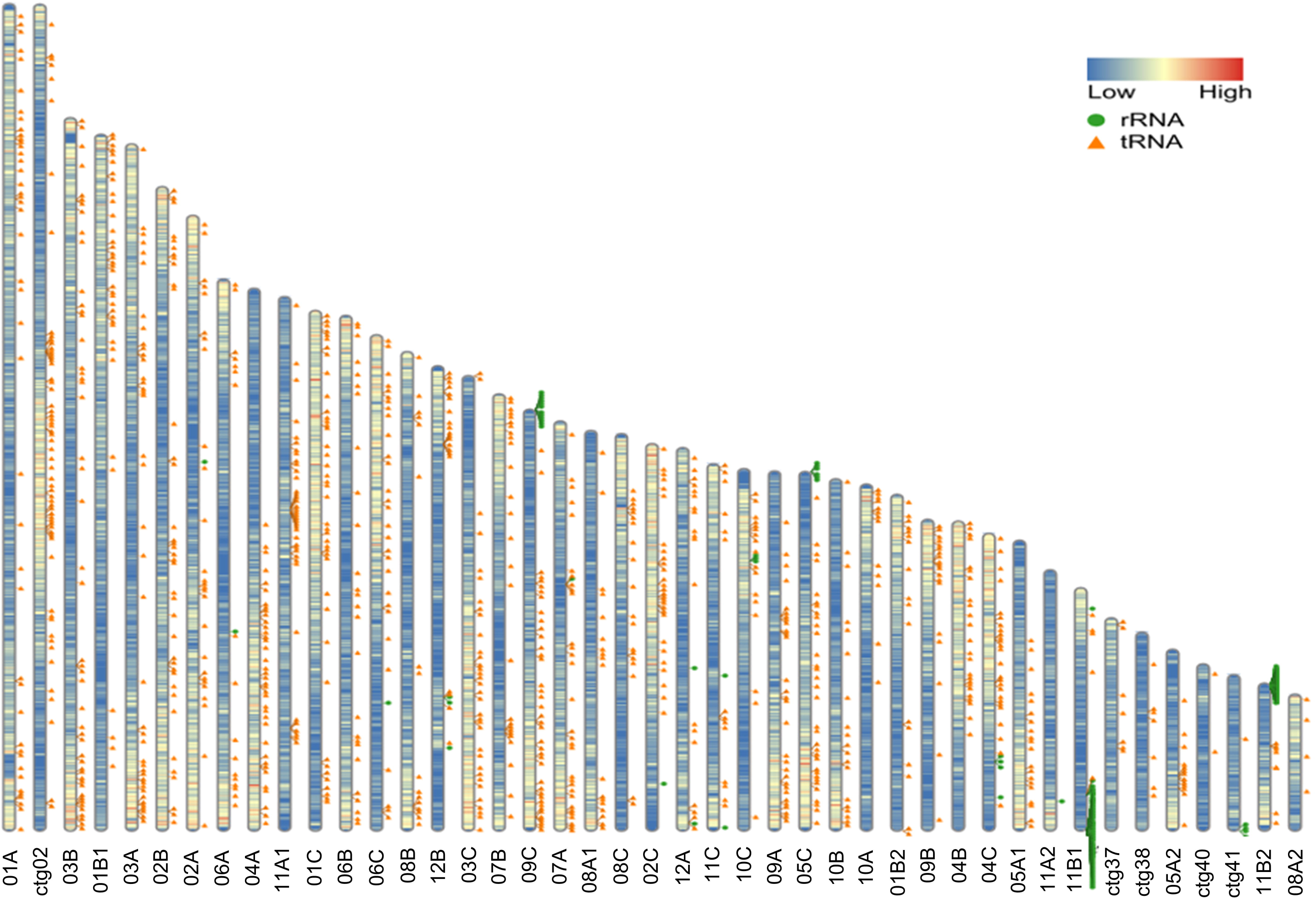
RIdeogram plot to visualize genome-wide gene density, tRNA position, and rRNA position across the 43 assembled contigs of *B. tulda*. The RIdeogram software was used for visualizing the genome features [51].

### Phylogenetic analysis

To determine the evolutionary relationship of *B. tulda* with other plants, twelve bamboo species (*Ampelocalamus luodianensis*, *Bonia amplexicaulis*, *Dendrocalamus latiflorus*, *D. sinicus*, *Guadua angustifolia*, *Hsuehochloa calcarean*, *Melocanna baccifera*, *Olyra latifolia*, *Otatea* glauca, *P.* edulis, *Raddia guianensis*, and *Rhipidocladum racemiflorum*) along with *Z. mays* and *Musa acuminata* (banana), were used to construct a species tree. OrthoFinder [48] was applied to the protein sequences of *B. tulda*, the twelve sequenced bamboos, and the two outgroups, *Z. mays* and *M. acuminata*. From the OrthoFinder analysis 202 orthogroups were obtained which were used for constructing the species tree (Fig. 6).

**Fig 6.**
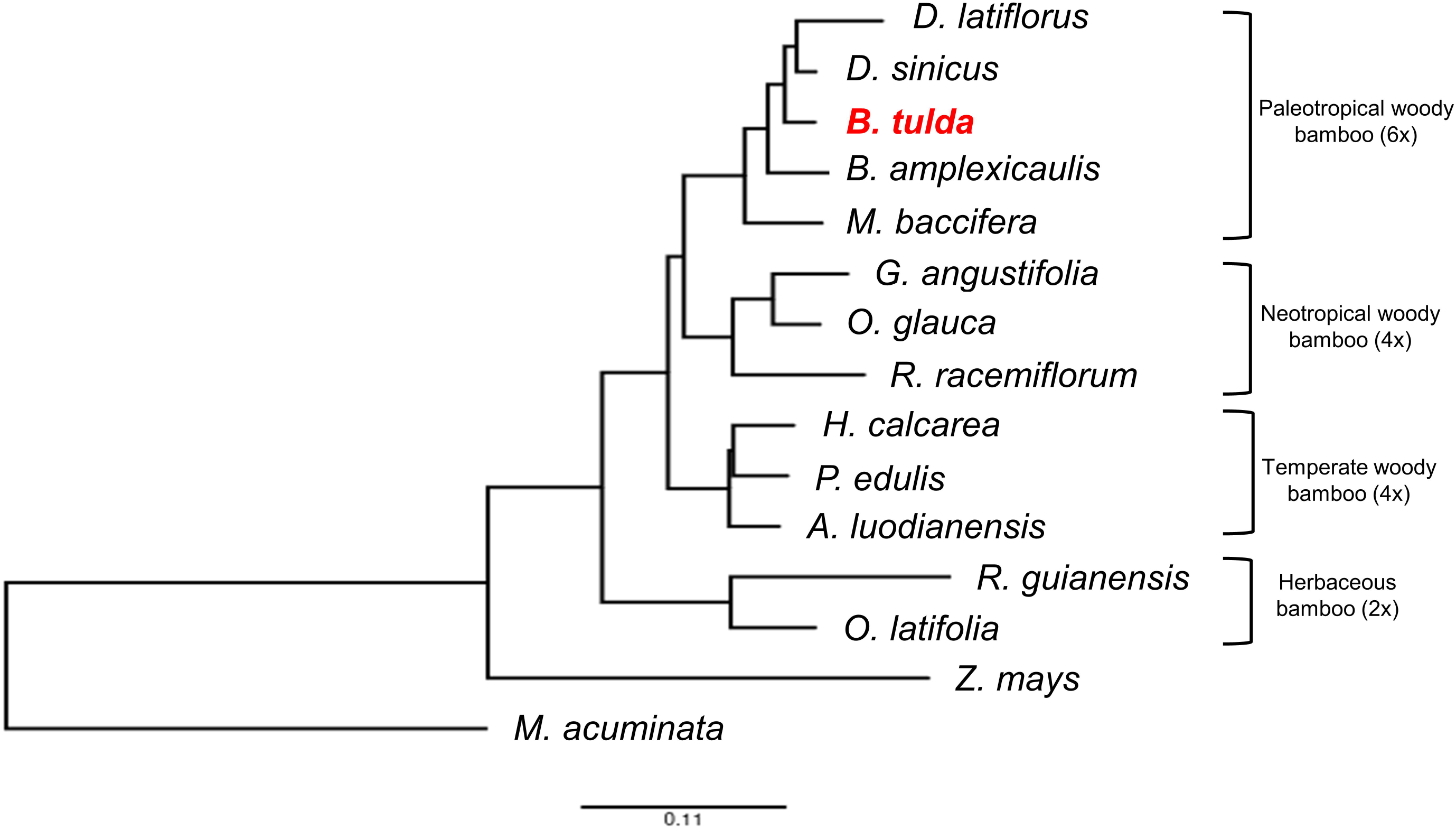
Species tree of *B. tulda* and other sequenced bamboos, with *Z. mays* and *M. acuminata* used as outgroups.

### Data Records

The raw PacBio genome sequencing data and raw Illumina RNA-Seq data were deposited in the EMBL-EBI European Nucleotide Archive (ENA) database under project accession number PRJEB91656, with sample accession numbers ERS25169745 for PacBio sequencing data and ERS25169746 for Illumina sequencing data. The genome assembly has also been submitted in the EMBL-EBI ENA database and will soon be made publicly available.

### Technical validation

#### Quality assessment of high molecular weight genomic DNA, libraries and sequence data

HMW genomic DNA was extracted to subject it for sequencing in the PacBio platform. The quantity and quality of the extracted DNA was examined by qubit fluorometer, and agarose gel (1%) electrophoresis (Fig. S1a). After ensuring required quality, three SMRTbell libraries were constructed from the genomic DNA, with relative fluorescence units (RFU) peaks obtained at 12,567 bp (DNA1), 11,447 bp (DNA2), and 11,307 bp (DNA3) (Figs. S1b, c, d). Each of these 3 libraries were sequenced in the PacBio Sequel IIe platform, while libraries DNA1 and DNA2 were sequenced in the PacBio Revio platform. The mean read lengths obtained in the Sequel IIe platform were 12,472 bp (DNA1), 12,524 bp (DNA2), and 12,864 bp (DNA3), while in the Revio platform were 10,033 bp (DNA1) and 13,155 bp (DNA2) (Fig. S2).

#### Assessment of genome assembly and annotation completeness

The completeness of the assembled genome was examined in BUSCO v5.8.0 [49] against the Embryophyte (n = 425) and Viridiplantae (n = 1614) datasets. BUSCO analysis demonstrated 99.0% complete, 4% fragmented, and 0% missing BUSCO groups against Embryophyte, and 98.0% complete, 3% fragmented, and 2% missing BUSCO groups against Viridiplantae (Fig. 3).

## Code availability

All softwares and pipeline utilized in this study for data analyses were implemented in full compliance with the manuals and protocols described by the respective published bioinformatics tools. Details on the software versions and parameters are outlined in the Methods section. In cases where specific parameters are not specified, default settings were applied. No custom programming or coding was employed.

## Acknowledgement

The research results reported in this paper are funded by the Alexander von Humboldt Foundation, Germany through ‘Research Group Linkage Program’. SK and SB acknowledge JRF fellowships from UGC, India, with NTA Ref. no.: 231610225668 and 211610047259 respectively. SD acknowledges CSIR Project (Grant no.: 38(1493)/19/EMR-II). We thank Prof. J. Dolezel for providing us the seeds of internal reference plants for genome size estimation. We thank Subhayan Paul, Institute of Health Sciences, Presidency University and Rajesh Saha, Bio-Rad Laboratories for providing technical support in operating the flow cytometer machine.

## Author contributions

MD and AB conceived the study, acquired funding and designed experimental plan. SK, SD, MB, and SB were involved in collection of plant samples, performing FACS experiments for genome size estimation, and isolating high molecular weight genomic DNA for high-throughput sequencing. SK and OR conducted all *in silico* experiments on genome data analysis. SK wrote the manuscript, and MD, AB and OR edited it. All authors read and approved the final manuscript.

## Competing interests

The authors declare no competing interests.

## Notes

https://www.ebi.ac.uk/ena/browser/view/PRJEB91656

### Competing Interest Statement

The authors have declared no competing interest.

