## Supplemental Figures for "Genome sequencing, *de novo* assembly and annotation of the commercially important bamboo, *Bambusa tulda* Roxb"

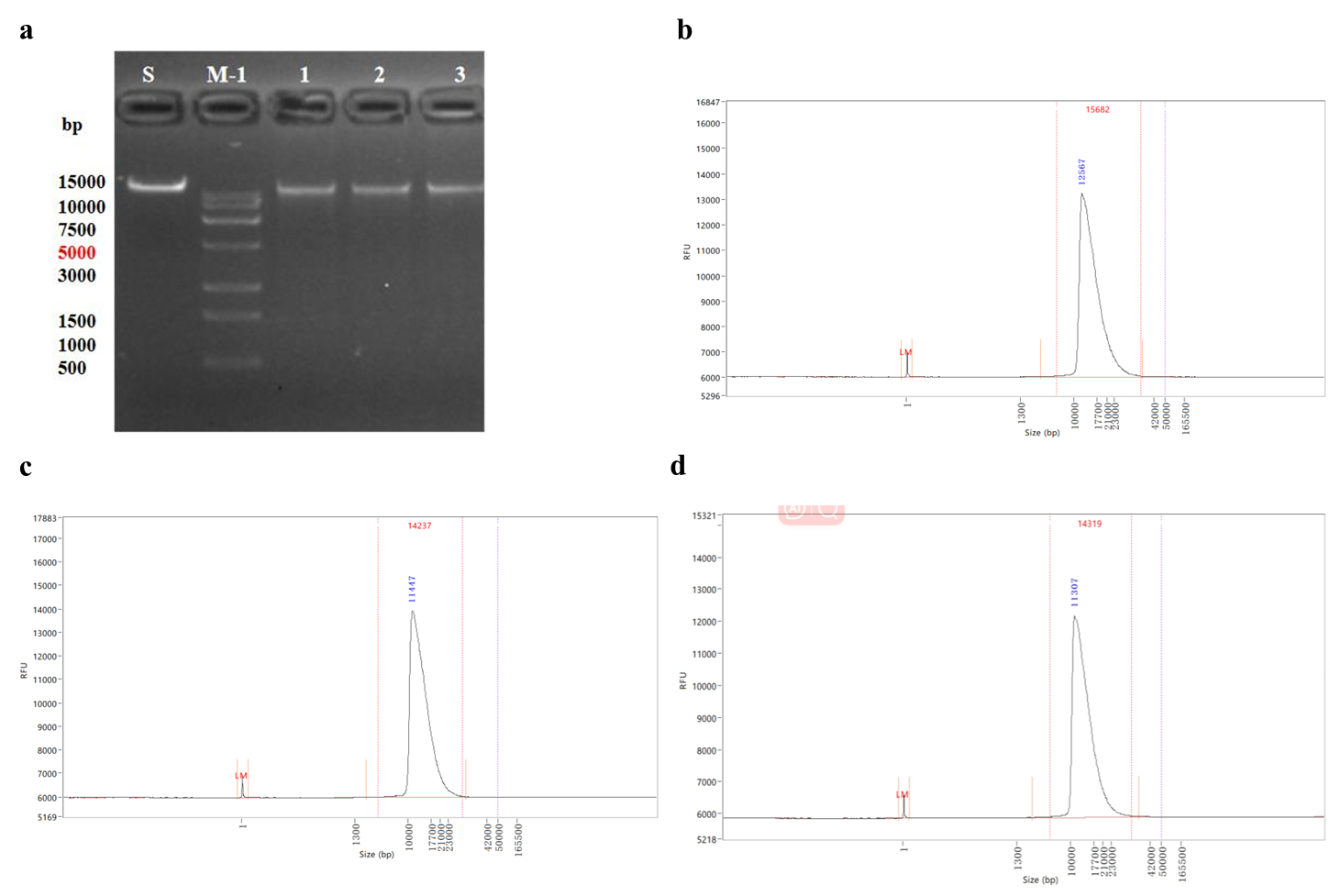
**Fig. S1.** (a) Quality assessment of the extracted HMW genomic DNA using 1% agarose gel electrophoresis. S denotes standard sample (50 ng), M-1 denotes trans 15k plus DNA ladder, 1-3: Three DNA samples diluted five times and loaded 1 μl. Bioanalyzer analyses of the three genomic DNAs used for sequencing in the PacBio platform; Library QC result for (b) DNA1, (c) DNA2, and (d) DNA3.

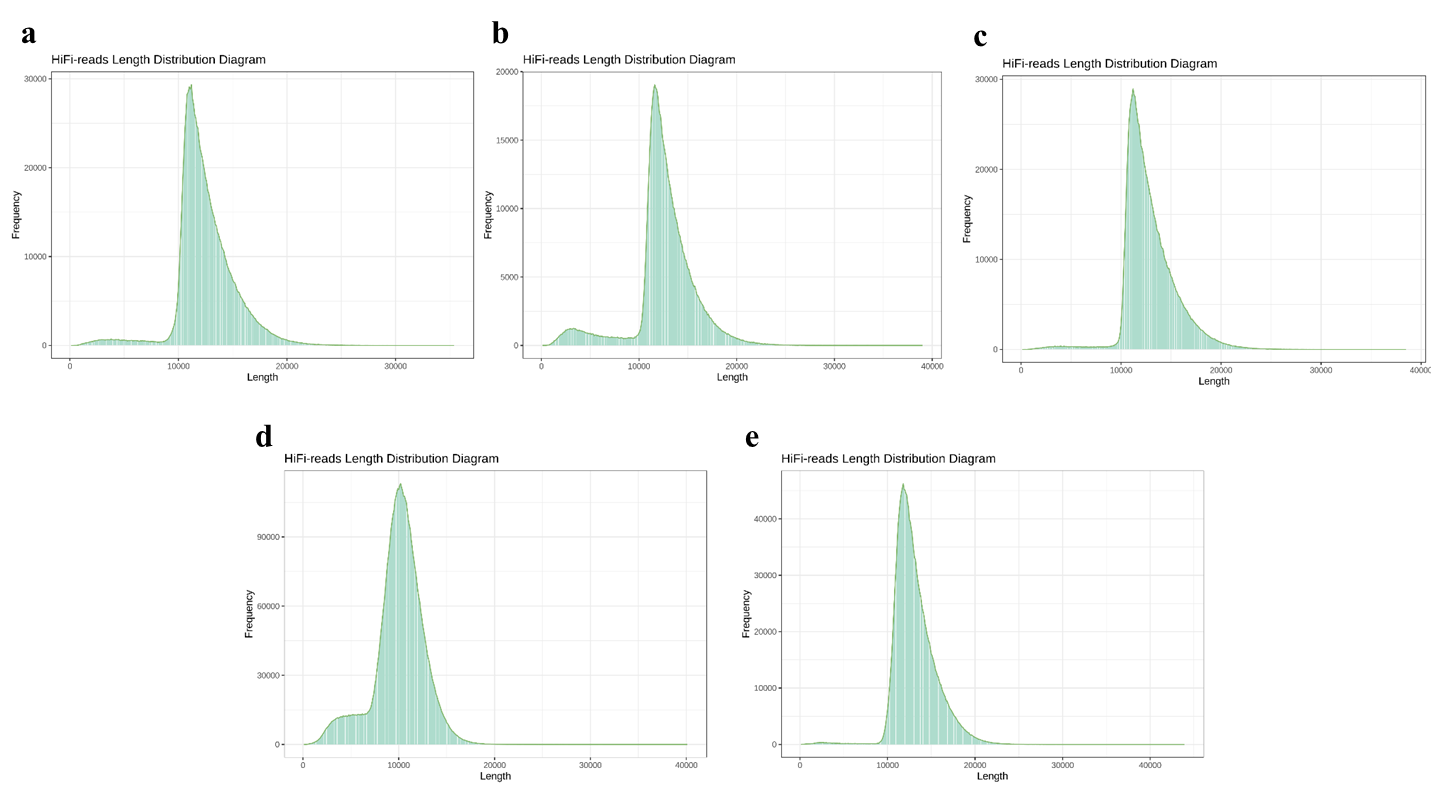

**Fig. S2.** Sequencing QC result for (a) DNA1, (b) DNA2, (c) DNA3 in the PacBio Sequel IIe platform, and (d) DNA1, (e) DNA2, in the PacBio Revio platform showing the mean read lengths.

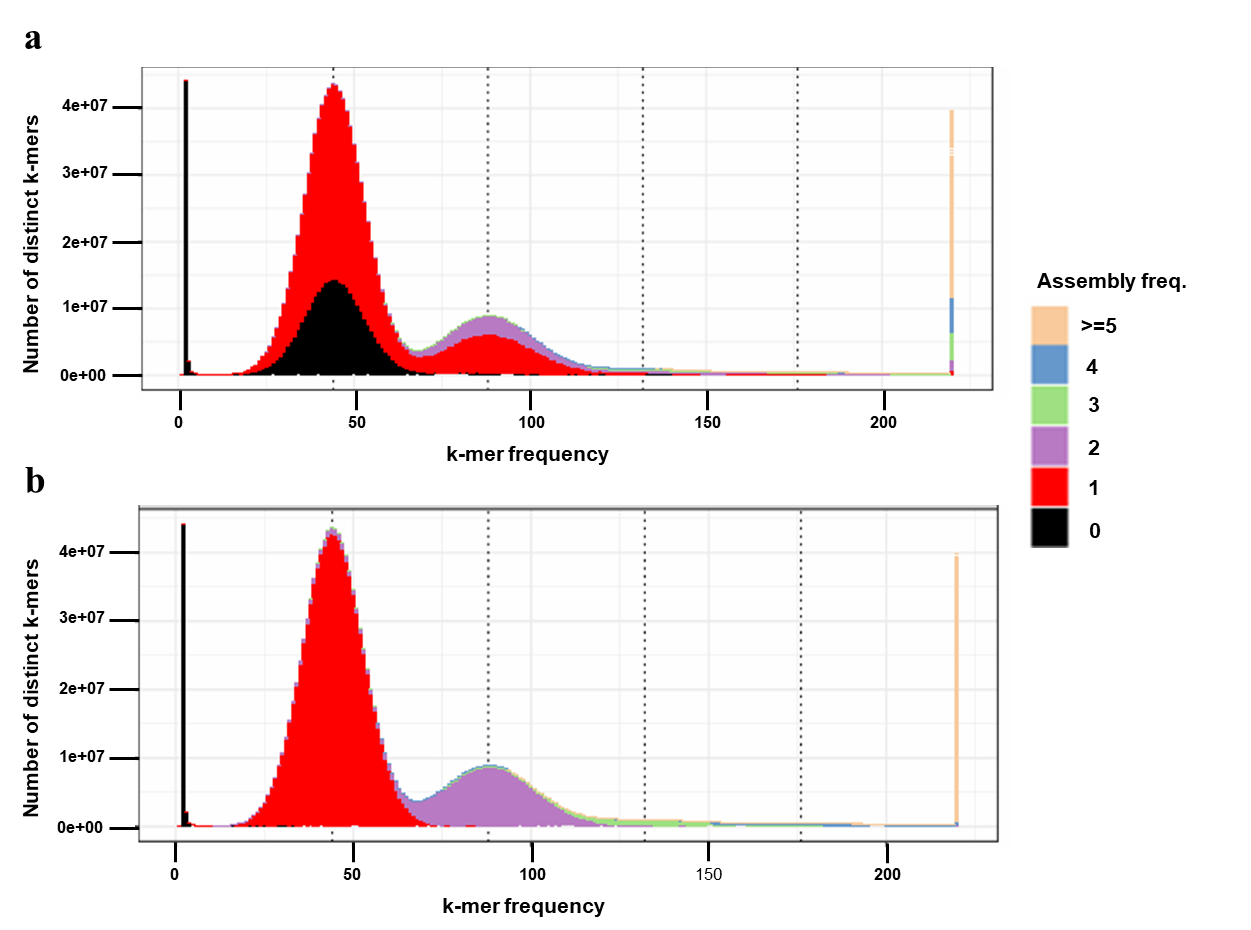

**Fig. S3.** *K*-mer comparison of genome assembly. (a) *K*-mer frequency spectra from reads to the primary contig assembly with only one haplotype. Majority of the *k*-mers occur only once (red) and 50% of the *k*-mers from the first peak are missing (black), denoting the haplotype which is not represented in this spectrum. (b) *K*-mer frequency spectra in the assembled contigs of both the haplotypes. The *k*-mers from the first peak occur once (red), from the second peak twice (purple), third peak thrice (green), and so on, in the assembly.
